# The Circadian Phase Response Curve to Light: Conservation across Seasons and Anchorage to Sunset

**DOI:** 10.1101/2021.04.08.439018

**Authors:** Hannah K. Dollish, Sevag Kaladchibachi, David C. Negelspach, Fabian-Xosé Fernandez

**Author notes:** Co-primary authors (Hannah K. Dollish, Sevag Kaladchibachi).

## Abstract

Phototherapy outcomes are dependent on the precision by which the magnitude and direction of light’s circadian phase-shifting effects can be predicted. Such predictions are largely based on photic phase-response curves (PRCs) generated from animals housed under seasonally agnostic equatorial photoperiods with alternating 12-hour segments of light and darkness. Most of the human population, however, lives at northerly latitudes (20-50° N of the Equator) where seasonal variations in the light-dark schedule are pronounced. Here, we address this disconnect by constructing the first high-resolution seasonal atlas for light-induced circadian phase-resetting. Testing the light responses of nearly 4,000 *Drosophila* at 120 timepoints across 5 seasonally relevant photoperiods (i.e., LD 8:16, 10:14, 12:12, 14:10, and 16:8; 24 hourly intervals surveyed in each), we determined that many aspects of the circadian PRC waveform are conserved with increasing daylength. Surprisingly though, irrespective of LD schedule, the start of the PRCs always remained anchored to the timing of subjective sunset, creating a differential overlap of the advance zone with the morning hours after subjective sunrise that was maximized under summer photoperiods and minimized under winter photoperiods. These data suggest that circadian photosensitivity is effectively extinguished by the early winter morning and out of optimal phase alignment with the wake schedules of many individuals (which revolve around the timing of sunrise regardless of season). They raise the possibility that phototherapy protocols for conditions such as seasonal depression might be improved with programmed light exposure during the final hours of sleep.

## 1. INTRODUCTION

The brain’s circadian pacemaker integrates information about ambient light and darkness to calibrate endogenous rhythms of physiology along a schedule that matches the 24-hour schedule set by the Earth’s rotation.^1,2^ This biological adherence enables animals to occupy a temporal niche phase-locked in alignment with the day (diurnal), night (nocturnal), or the temporal poles stationed between them at dawn and dusk (crepuscular), thereby organizing *sui generis* behaviors related to sleep, feeding, and reproduction to times of the solar cycle optimal for survival.^3^ The pacemaker not only evolved to anticipate daily recurring episodes of consolidated light exposure followed by darkness but also to annual variations in the length of the day *relative to the night*.^4^ Owing to the Earth’s 23.45° axial tilt and orbit around the Sun, the ratio of day versus night changes systematically across the year, with the degree of change dependent on latitudinal (angular) distancing north or south from the equator. When the boundaries separating the day from the night are moved by shortening or extending the duration of the daily light period, the pacemaker will continue to direct 24-h rhythms while repositioning the onset and offset of many physiological indices to the new transitions marking sunrise and sunset.^4-7^ The most conspicuous of these indices is the onset and offset of nightly melatonin secretion, which imprints a biological representation of daylength animals use to make forward-looking seasonal adaptations (e.g., gonadal regression, hibernation) to predictable environmental challenges introduced at times of year when the number of daylight/night hours becomes significantly skewed (e.g., severe decreases in temperature and food availability in winter when the night is extended at the expense of the day).^7-9^

While much is known about the independent mechanisms that subserve the pacemaker’s operation as a circadian clock versus photoperiodic seasonal timer^10-23^, comparably less is understood about the intersection of these two areas. Per current knowledge, the pacemaker’s phase responses to acute light exposure change systematically across the subjective night in the service of photoentrainment to an equinoctial 12:12 light-dark (LD) cycle.^24^ Here, light administration in the first half of the evening will produce phase *delays* in behavioral-physiological rhythms, whereas administration in the latter half will produce phase *advances* (light stimulation during the subjective day tends to provoke little phase movement either way).^24,25^ It is assumed that these responses stem from the pacemaker’s differing interpretations of light exposure in the several hours subsequent to dusk as opposed to the several hours preceding dawn. In the former, any incidence of light telegraphs an extension of the sunset requiring delays in activity offset that will maximize a diurnal animal’s contact with daylight; in the latter, light signals an earlier sunrise requiring advances in activity onset that will enable a diurnal animal to awake in alignment with the start of the morning (reciprocal relationships hold for nocturnal animals). When graphing these responses across hourly intervals, it is customary to plot the magnitude of delays with negative numbers and advances with positive numbers.^26^ The resulting visual that emerges—termed the circadian phase response curve (PRC)—takes on the appearance of a sigmoidal-like wave that zeroes out near the scheduled transitions of the light-dark cycle.^24-26^

The circadian PRC to light has been predominantly studied in model organisms (e.g., mice and flies) entrained to an equatorial photoperiod.^27,28^ To our knowledge, a comprehensive PRC atlas detailing how patterns of circadian photosensitivity change as a function of daylength across several seasonally relevant photoperiods has never been constructed. Such an atlas has the potential to open a discussion about whether current practice guidelines for phototherapy require revision to accommodate differences in the PRC that might develop as a function of time-of-year and geographical residence. The success rate for phototherapy in conditions such as seasonal affective disorder is heavily dependent on the precision by which the PRC can predict the magnitude and direction of light’s circadian phase-shifting effects.^29-31^ The variability of treatment success observed in these clinical areas^32-34^ might be due (in some measure) to the fact that phototherapy protocols have been standardized according to PRCs generated within seasonally agnostic LD schedules comprised of light and darkness available in equal measure for 12 hours.

A symmetrical split between day and night is observed at the equator (0°) where it remains relatively fixed regardless of season. However, moving away latitudinally from 0°, photoperiods become more dynamic in their seasonal variation. Although many people live at or around the equator (+/- 5°), data from the NASA Socioeconomic Data and Applications Center (SEDAC) indicate that global population density is highest at 30° N/S, where summer and winter photoperiods range from LD 14:10 to 10:14, respectively. Beyond 30° N/S, seasonal variations in photoperiod are even more dynamic; heavily populated regions at this latitude such as Toronto, New York City, and Beijing (40° N) rapidly fluctuate in daylength as the year progresses, going from LD 16:8 in summer to 8:16 in winter and toggling back and forth from these extremes in a matter of months. The geographical distribution of the human population with respect to the equator suggests that the circadian pacemaker of most individuals—at most times of year—is under the influence of photoperiods that deviate significantly from LD 12:12.

In the current study, we constructed high-resolution PRCs for 5 photoperiods exhibiting incremental changes in the ratio of light and darkness (LD 8:16, 10:14, 12:12, 14:10, and 16:8). This range provided an accurate representation of winter → summer LD schedules, allowing us to determine how circadian photosensitivity might change across the seasons in geographical areas where the share of the global population is highest (20-50° N/S, **Figure 1**). We outline the results below and discuss their clinical relevance with respect to the PRC waveform and its localization relative to the timing of sunset and sunrise.

**Figure 1.**
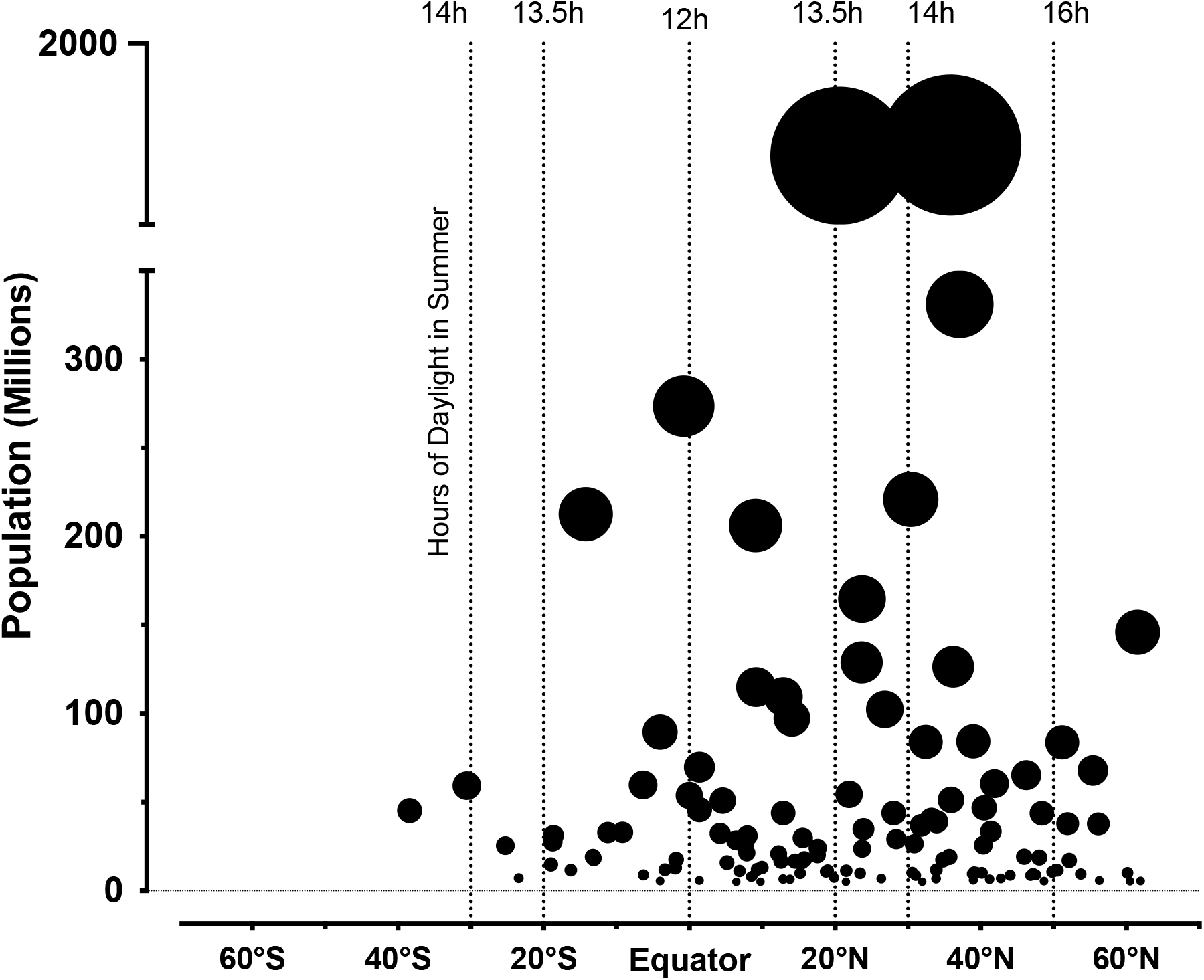
Countries in the world are plotted as individual points with respect to their central latitude (°N/S) and population (in the millions). The size of each point is scaled to population level (referenced from United Nations Population Division). Dotted lines intersecting each latitude mark the seasonal swings in daylight associated with countries located at each deviation from the Earth’s Equator. The number of daylight hours during summer (and the inverse in winter) increases moving away from this midline. The majority of the human population lives between 20-50° N, where up to 8-hour swings in the photoperiod occur every 6 months.

## 2. METHODS

*D. Ananassae* (melanogaster species subgroup) were derived from an isofemale line maintained at the *Drosophila Species Stock Center* (DSSC) at Cornell University’s College of Agriculture and Life Science (stock # 14024-0371.16). Stocks were reared at 25°C in DigiTherm® incubators (Tritech Research, Inc., Los Angeles, CA) and entrained to a 12:12 LD cycle (broad-spectrum light source: 4 watt cold-cathode fluorescent light tube with step-up inverter, freely mounted with no fixture, illuminance at rack level = 887.7 lux; Tritech model DT2-LB-F12IN/CIRC-L-INV; lights-on at 07.00 h, MST). The stocks were transferred daily to generate a steady supply of offspring. For phase-shifting experiments, female flies were selected as late-stage, “pharate-adult” pupae, moved onto fresh food, and housed in groups of 5 to 6. A few days post-eclosion, animals were singly housed in Pyrex glass chambers (5 mm outside diameter, 65 mm long) containing a solid aqueous plug of food medium (4.7% sucrose, 1.9% agar; MoorAgar, Inc., Rocklin, CA) on one end and a cotton fitting on the other, and loaded into Trikinetics DAM2 Drosophila Activity Monitors (TriKinetics, Inc., Waltham, MA). The monitors were situated within climate-controlled vivariums identical to the ones used in colony management but with the photoperiod maintained, extended, or retracted to one of five possible seasonal schedules: **LD 8:16** (lights-on at 0.700; lights-off at 15.00), **LD 10:14** (lights-on at 07.00; lights-off at 17.00), **LD 12:12** (lights-on at 07.00; lights-off at 19.00), **LD 14:10** (lights-on at 07.00; lights-off at 21.00), or **LD 16:8** (lights-on at 07.00; lights-off at 23.00). Throughout the experiment, the flies’ motion was independently tracked by cross-sectioned infrared beams, which transmitted movement information over modem/USB to a computer acquisition software (DAMSystem-3) every 30 seconds.

An Aschoff Type II paradigm was used to quantify the effects of light exposure on phase resetting of the flies’ locomotor activity rhythms. Animals were entrained to the LD schedule to which they were introduced for 3 days. After lights-off on the last day of the entrainment period, separate cohorts received a single 15-min pulse at one of the hourly intervals spanning the next 24-h cycle (i.e., 24 independent groups tested per seasonal schedule; 120 groups in total). This was accomplished by software-controlled activation of the house lamp (887.7 lx, white fluorescent light; Tritech Research, DeviceCom3™). Post-pulse, animals were left to free-run in constant darkness (DD) for 4-5 days.

Actogram plots reflecting the daily activity profile for each fly in a given treatment group were created by binning raw 30-sec time series data of individual *D. ananassae*. Phase shifts of behavior were calculated for each fly (one fly = one 5×65 mm tube) by determining the horizontal distance between regression lines fitted through software-called activity onsets 2 days prior and 2-4 days after light administration (ClockLab Analysis Version 6, Actimetrics, Wilmette, IL). Initial experiments assessing the flies’ activity profile within each photoperiod showed that the activity onsets of *D. ananassae* were always phase-locked to the timing of lights-on in each seasonal schedule (i.e., 07.00 h). Post-pulse, transients were observed for a day, but the flies’ behavioral rhythms stably reset by the second DD cycle (hence the start of the regression here). To correct for phase movements that might simply accompany transitions from LD to DD in each seasonal condition, a control group was transferred into DD without any light treatment. Net calculations of onset shifts were normalized for the effects of removing the LD cycle. For analyses, one-sample *t*-tests were used to determine whether the net phase shift produced by light exposure at a given hour for a given seasonal schedule was greater than zero; this value indicates no phase movement. Where appropriate, one-way or two-way ANOVAs with Tukey’s post hoc correction were used to assess seasonal variations in phase-shift magnitude across select intervals encompassing the delay and advance zones. Significance was set at *p* = 0.05. In total, 3,934 animals were independently evaluated to measure the effects of seasonality on light-induced circadian resetting, while another 160 were used to examine seasonal activity profiles. Please note that the phase-shift data for the hourly intervals between ZT13 to ZT23 in the 12:12 LD condition are reproduced from Kaladchibachi et al., 2018^35^, which reports on a battery of experiments largely contemporaneous to the ones discussed in the current manuscript.

## 3. RESULTS

Flies maintained robust 24-h patterns of locomotion under each photoperiod tested. In all cases, activity profiles were marked with morning and evening peaks that were coincident with the timing of lights-on and lights-off in the LD schedule (**Figure 2A**). While morning peaks of activity gave way to similar declining rates of movement leading into midday, evening peaks scaled with the number of hours available in the photophase; the longer the daylength, the more that evening bouts of activity encroached into the photophase, encompassing a span of about 6 hours when animals were housed in a LD 16:8 (**Figure 2A-B**). Morning anticipatory activity was not a general feature observed in the photoperiods that were tested, though such patterns emerged after a few days of housing under a high-winter 8:16 LD schedule (**Figure 2B**). Upon complete removal of the LD cycle and entry into DD, activity onsets were increasingly phase-delayed with respect to the timing of lights-on (dawn) in the previous photoperiod; the longer the daylength in the previous schedule, the larger the delay that was observed when the LD cycle was eliminated (**Figure 2C**; *F*_4,155_ = 726.3, *p* < 0.0001). The scaling of this phase movement is consistent with the photoperiodic aftereffects noted by Pittendrigh in the *Drosophila* eclosion rhythm upon steady-state release into DD from a range of seasonally-relevant photoperiods^36^. The irradiance spectrum for the broadspectrum light source used to generate the photoperiods and construct the PRCs described below is provided in **Figure 2D**.

**Figure 2.**
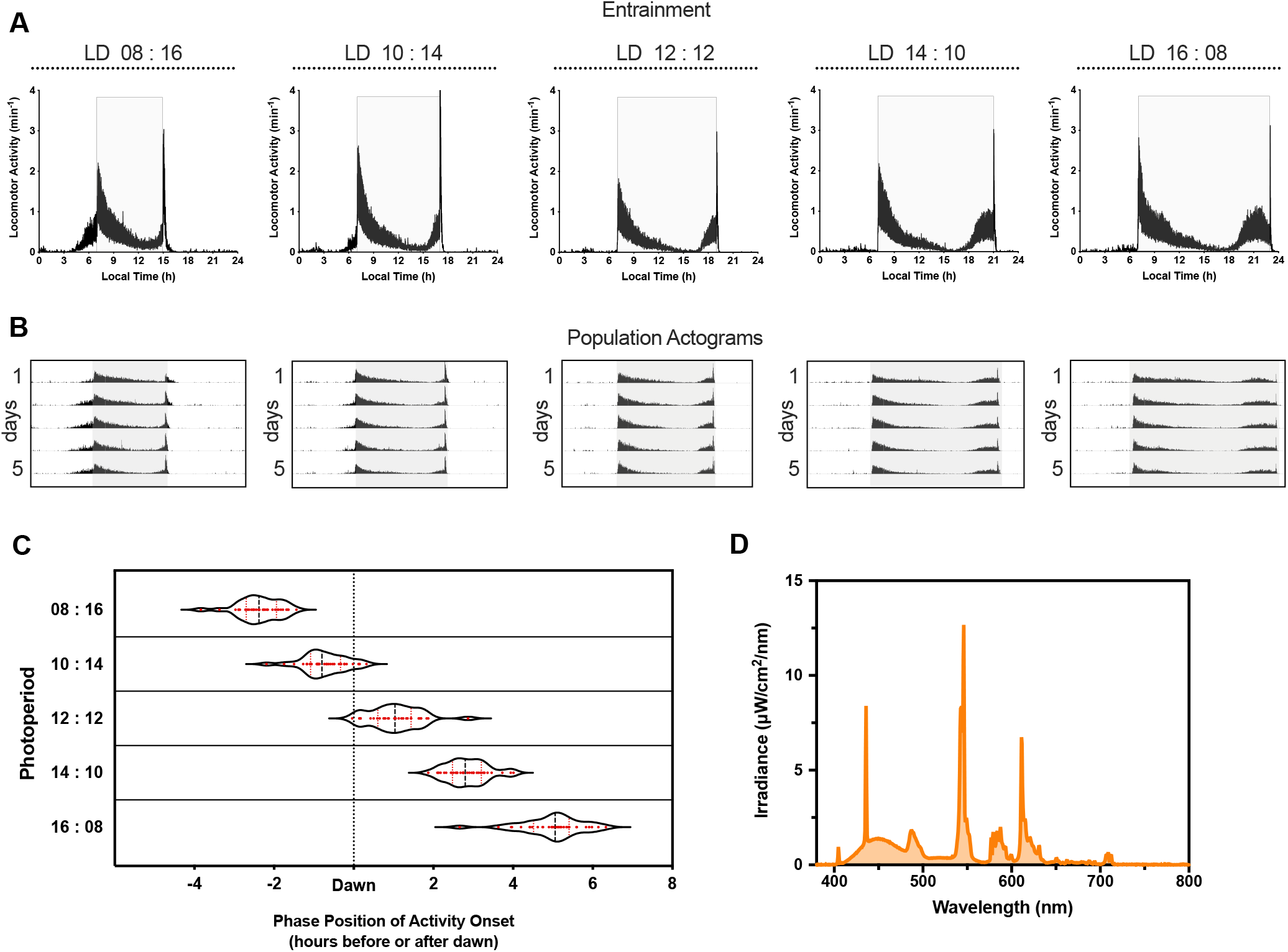
Entrainment to seasonal photoperiods. (**A**) Eduction plots showing the 24-h activity profiles (min^-1^) for flies maintained under the various photoperiods. The LD cycles are labeled overhead, left to right, with the light phase in each bracketed in light gray. Activity profiles were created by binning raw 60-sec time series data of individual flies, averaging these data across animals, and then averaging across each corresponding 60-sec epoch of each day under study (entrainment was monitored for 5 days). (**B**) Population actograms for flies housed under each photoperiod are provided, situated below each of the activity profiles. Black bars indicate 30-sec epochs where the animals registered breaks in the infrared-tracking beams, smoothed over each two minutes of recording. Data are vertically aligned, such that one 24-h day of movement is shown per line, with successive days appearing one below the other. The portion of the LD schedule where the lights are on is highlighted in grey (local time of lights-on in each photoperiod = LD 08:16, 07.00-15.00; LD 10:14, 07.00-17.00; LD 12:12, 07.00-19.00; LD 14:10, 07.00-21.00; and LD 16:08, 07.00-23.00). (**C**) Violin plots show the magnitude of the phase-shift in each animal’s locomotor activity onset calculated 2 days after removal of the light-dark cycle from each photoperiod (1 point = 1 fly). Data are plotted with respect to the timing of lights-on in the previous schedule (Dawn). The overall distribution of the data points (red) is reflected by changes in width that occur along each plot. The median value and quartiles of the samples are marked with a thick fragmented line and thinner red-dotted lines, respectively. (**D**) Irradiance spectrum for the broadspectrum light source used in the study. Illustrated within are the dominant wavelengths produced by the emission.

Raw data from the PRC experiments are reported in five scatterplots visualizing phase changes in the locomotor activity rhythm for individual animals exposed to a 15-min pulse at one of the hourly intervals of the 24-h day (**Figure 3**; descriptive statistics are available in **Supplementary Tables 1-5**). These data were averaged at each timepoint to generate high-resolution PRCs that illustrated how the circadian pacemaker’s sensitivity changed as a function of seasonal photohistory (**Figure 4**). In theory, seasonal differences in the PRC might manifest as differences in the fraction of the circadian cycle that light-induced resetting of any kind occurs. If this fraction remains constant, there might still be changes in the amplitude of the delay or advance zone (i.e., scaling of the maximal shifts achievable by saturating light exposure), retractions or expansions in the widths of these regions (i.e., changes in the proportion of the PRC where light exposure will elicit a delay versus an advance shift), or changes in the positioning of the delay and advance zones with respect to the timing of lights-on (dawn) and lights-off (dusk) in the LD schedule.

**Figure 3.**
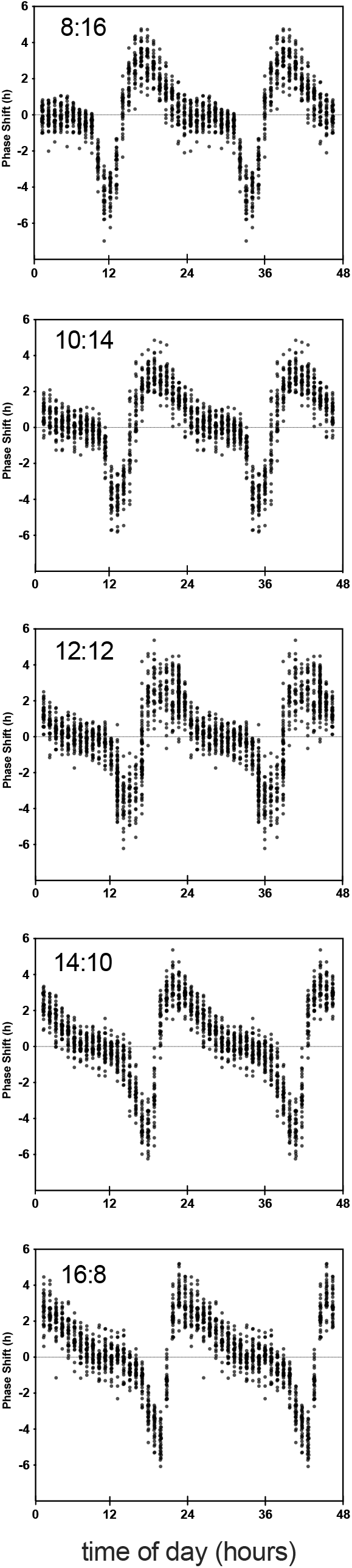
Phase-shifting responses across the seasons: Raw data. Phase shifts of locomotor activity (h) are quantified for individual flies (n = 3,934) stimulated with a 15-min pulse (white fluorescent light, 887.7 lx) at one of the hourly intervals associated with the 24-h cycle starting after the last day of entrainment to each LD schedule (post-pulse, animals were maintained in DD). Data collected from animals housed under each photoperiod are independently graphed and labeled from top-to-bottom. The phase shift observed in each fly’s activity rhythm (1 black circle = 1 animal) is shown in scatterplot with respect to the timing of the pulse. X-axes start at ZT0, which reflects the timing of lights-on (dawn) in the previous LD schedule. Data are double-plotted for ease of visualization.

**Figure 4.**
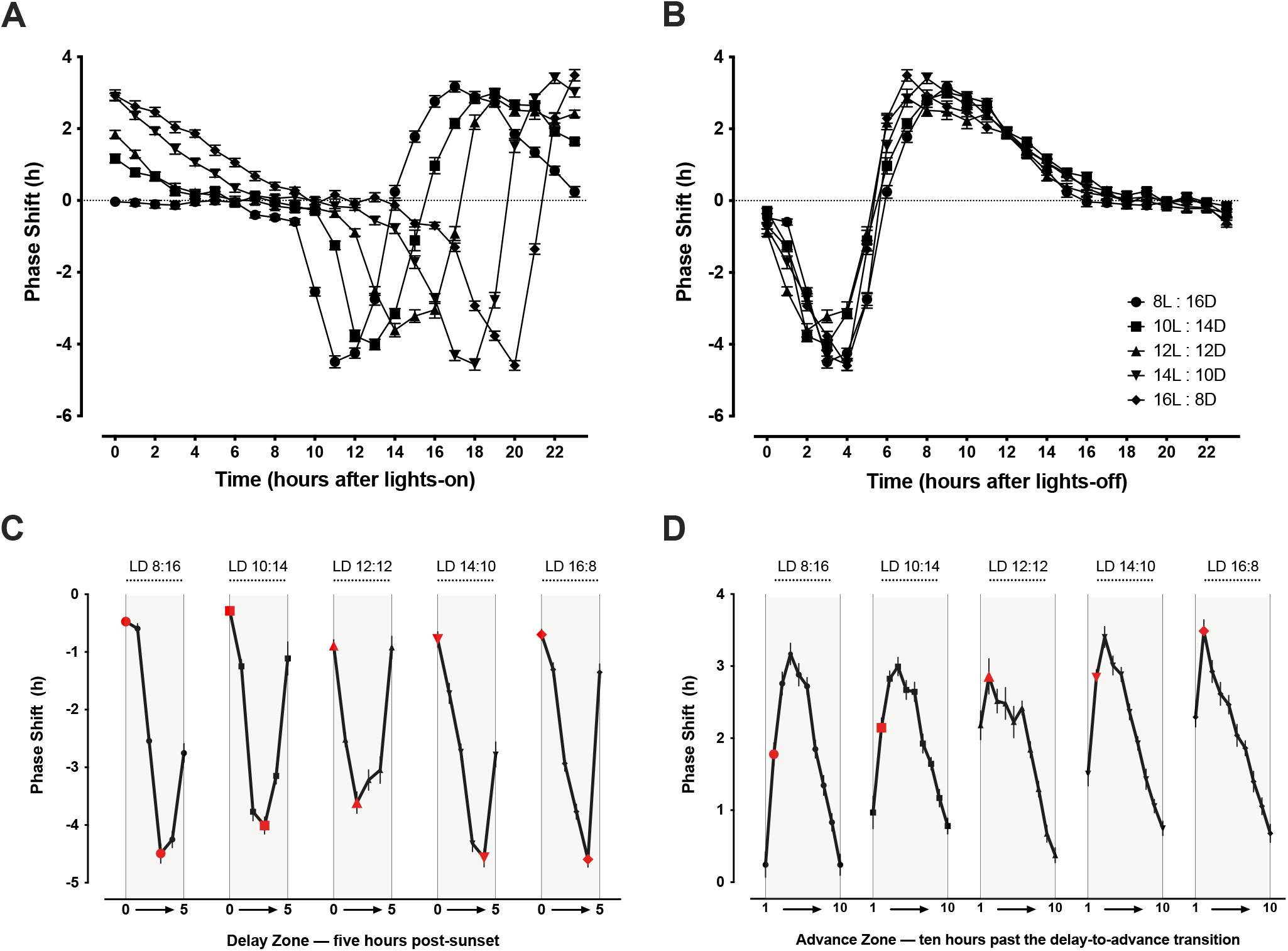
Phase-shifting responses across the seasons: Averaged data. (**A-B**) The PRCs generated after entrainment to each photoperiod are co-plotted with respect to the timing of either lights-on (subjective sunrise, **A**) or lights-off (subjective sunset, **B**) in the previous LD schedule. Independent of photoperiod, PRCs are anchored to the timing of lights-off. (**C-D**) The parts of the PRC corresponding to the delay (**C**) and advance (**D**) zones are isolated for each photoperiod (labeled left-to-right) to visualize similarities in the trajectories of circadian photosensitivity occurring along each curve. Red points highlight hourly stretches where these trajectories were slightly, but statistically different. Inflections of photosensitivity (moving from the amplitude of the delay → advance zone) were sharper for longer photoperiods with shorter nights.

In the midst of all these possibilities, we found very targeted effects of seasonality on the circadian PRC to light. Primarily, this involved phase-positioning of the entire curve relative to the transition borders of the LD cycle. Irrespective of season, photic PRCs were always anchored to the timing of lights-off (subjective sunset) in the LD schedule, producing a staggered realignment of the advance zone relative to the timing of lights-on (subjective sunrise) (**Figure 4A-B**). The absolute widths of the delay and advance zones were conserved—comprising approximately 5 and 10 hours of the circadian cycle, respectively, along with a 9-hour deadzone (**Figure 4B-D**)—though *minor* photosensitivity differences were distinguishable across hourly intervals within each region. A two-way ANOVA for the phase-shift data within the first 5 hours of the scotophase, the interval marking the delay zone, indicated a photoperiod/seasonal effect (*F*_4,1007_= 9.471, *p* < 0.0001), circadian phase effect (*F*_5,1007_= 344.3, *p* < 0.0001), and an interaction between season and circadian phase (F_20,1007_ = 19.69, *p* < 0.0001) such that photosensitivity peaked later with summer-like photoperiods possessing ≥ 14 hours of daylight (**Figure 4C**). A two-way ANOVA for the phase-shift data within the 10-hour block of the circadian cycle aligned with the delay-to-advance transition (marking the advance zone) also indicated a photoperiod/seasonal effect (*F*_4,1576_ = 9.448, *p* < 0.0001), circadian phase effect (*F*_9,1576_ = 170.5, *p* < 0.0001), and an interaction between season and circadian phase (*F*_36,1576_ = 7.759, *p* < 0.0001) such that photosensitivity was greater within the first two hours of the advance zone under longer photoperiods with short nights (**Figure 4D**). The net effect of these changes was to create a sharper inflection in the waveform of the PRC as it transitioned from the most photosensitive area of the delay zone to the most photosensitive area of the advance zone (**Figure 4B**).

In one final analysis, we replotted phase-shift data relative to the timing of lights-on (subjective sunrise) in each photoperiod to quantify the number of hours within the first 8-hour segment of the morning/afternoon that remain circadian photosensitive by virtue of the morning’s overlap with the PRC advance zone (**Figure 5**). Overlap of the advance zone with the morning was minimized under short, winter-like photoperiods (e.g., LD 8:16 and 10:14) and accentuated under long, summer-like photoperiods (e.g., LD 10:14 and 16:8). At the seasonal poles, we found that the pacemaker’s photosensitivity was effectively extinguished by sunrise during high winter but robust throughout much of the first-half of the day in high summer.

**Figure 5.**
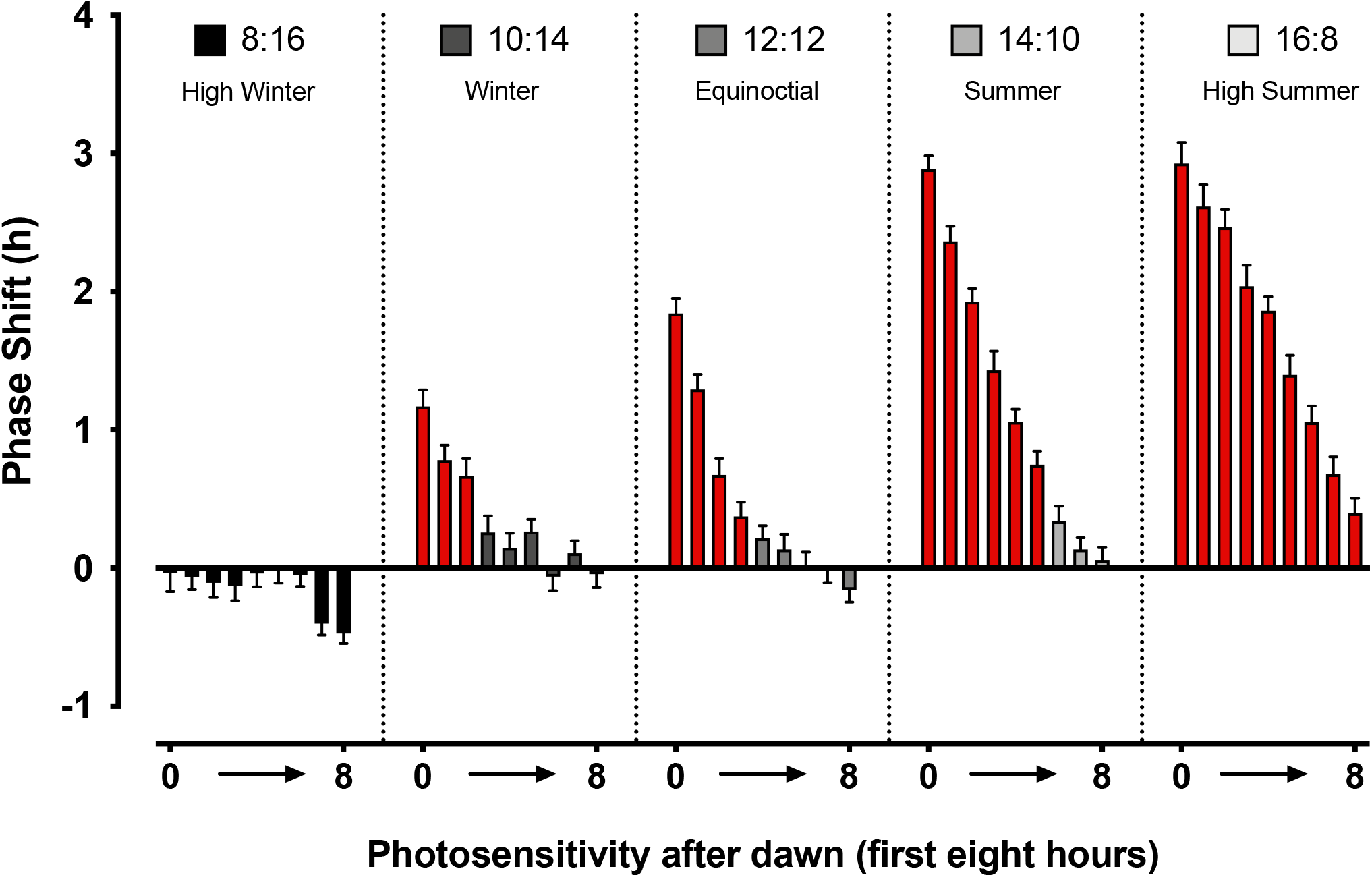
Phase-shifting responses across the seasons: Focus on the morning hours. The average phase-shift achieved with a 15-min pulse was replotted for each seasonal photoperiod (labeled left-to-right) and referenced to the hourly intervals of the circadian cycle coinciding with the first 8 hours of the morning/afternoon after lights-on (dawn). Red bars indicate times where light exposure produced a net phase-advance greater than zero (one-sample *t*-test, *p* < 0.05). Overlap of circadian photosensitivity with the morning/afternoon hours was maximized in summer photoperiods and minimized (or effectively absent) in winter photoperiods.

## 4. DISCUSSION

Here, we report development of the first seasonal atlas for light-induced phase resetting of circadian rhythms. After probing 120 timepoints across 5 seasonally relevant photoperiod schedules, we determined that many aspects of the circadian photic PRC are conserved with increasing daylength, including the widths and amplitudes of the delay and advance zones and the proportion of the curve comprised by each region. The single largest impact of seasonality on the PRC was to change the extent to which circadian photosensitivity in the advance zone extended into the morning hours after sunrise. By virtue of the PRCs’ anchor to sunset, this intersection was maximized under summer photoperiods and limited under winter photoperiods with ≥ 14 hours of darkness.

It is worth noting that there has been only one previous systematic attempt to understand how the pacemaker’s light responses are affected by photoperiod. In studies that were never formally published (but disseminated as part of a larger book chapter), Elliott and Pittendrigh found that the amplitude of the hamster PRC to light was significantly larger when animals were housed under a LD cycle with short days (LD 10:14) versus one with long days (LD 18:6)^37^. Though it was never directly articulated by the authors, examination of their plots suggests that both PRCs were pegged to the timing of lights-off (subjective sunset) in each photoperiod, with a resulting: 1. several-hour overhang of the advance zone with the morning/afternoon hours of the photophase in the LD 18:6 schedule and 2. absence of circadian photosensitivity in the morning hours of the winter-like LD 10:14 schedule (see Figure 5 of reference 37). These data, collected across two photoperiods probing 9 timepoints in each, are limited relative to the current seasonal atlas but are largely consistent with the findings we have made. Our two data sets might be further generalizable still in view of later experiments done by Elliott and his collaborators (Glickman and Gorman)^38^, who determined that the amplitude difference in the hamster PRC under short versus long days is nullified (at least at one circadian phase) when light exposure beyond ∼100 lux is used to generate phase responses; in other words, when ambient illumination commensurate with room lighting is used, the responses are equivalent.^38^ While seasonal differences are bound to exist in the pacemaker’s responses to dim versus bright light, or broadspectrum versus narrowband emission, or light delivered along other physical exposure dimensions, the current study establishes that the PRCs reflecting maximal responses across the seasons do not generally change in their waveform characteristics or their phase-positioning relative to sunset. All the aforementioned considered, these tenets are likely to be species generalizable and may be pertinent to the practice of phototherapy in humans.

Phototherapy guidelines set by the medical community and health insurance industry for conditions thought to have an underling circadian basis such as seasonal affective disorder (SAD) suggest timing light administration within the morning as close to waking as possible.^39^ For many individuals residing in the “human hemisphere” where population density is highest at 20-50° N, this circadian phase <u>may no longer coincide</u> with any remaining pacemaker light sensitivity during the winter months^40,41^ when SAD mood symptoms are at their worst. For those individuals, a tangible translation of our results to the clinic might entail the development of home-based healthcare provider technologies that can time light administration during sleep for the several hours preceding a person’s habitual waketime. Such strategies might afford better treatment efficacy and would almost certainly lead to better treatment compliance considering the well-known side effects (e.g., headache, eye strain, jitteriness) and adherence issues that accompany current insurance-covered “bright-light therapy” protocols.^42,43^

In summary, we have established a seasonal atlas for circadian phase-shifting and identified the advance zone as the major site of the PRC undergoing photoperiod-dependent realignments in positioning with respect to the LD cycle. Future experiments will be necessary to scrutinize other aspects of daylight and twilight that might add important nuances to how photic PRCs are influenced by seasonality. For instance, seasonal variations in the spectral composition of sunlight—and patterns of spectral change over the daytime—mark the photoperiod at all latitudes.^44^ Introduction of more natural “dynamic” lighting conditions to phase-shifting experiments, though rarely (if ever) practiced in the laboratory, might thus refine PRC waveforms and have special significance for identifying changes to photic PRCs constructed using narrowband and intermittent photic stimuli.^45,46^ Another future consideration involves the use of graded twilight simulations. Though virtually all phase-shifting studies to date have been conducted in the laboratory under square-wave LD cycles carrying abrupt (switch-like) transitions between lights-on and lights-off, inclusion of natural twilight progressions—over the course of 60-90 minutes—might signal (or aid in the communication of) other seasonal factors to the pacemaker that impact the shape of the PRC manifesting under different photoperiods.^47-49^ Lastly, the PRC’s changing alignment with the day and night suggests that the relationship between circadian phase-shifting and melatonin might change across the seasons. In the simplest example, during winter, when photic PRCs are completely compartmentalized to the night, melatonin might play more of a role as a natural zeitgeber and adjunct therapeutic agent alongside light in the timing of the wake-sleep cycle.^50-53^ All of these considerations underscore the need for characterization of the circadian pacemaker outside the typical contexts (e.g., equatorial LD schedules) in which it has been historically studied. Such insights have the potential to enhance the visibility of phototherapy in the medical and psychiatric research communities and improve treatment outcomes.

## Acknowledgements

We are indebted to the Velux Stiftung (Proj. No. 1360) for their financial support.

## Author Contributions

F. Fernandez developed the study concept, oversaw its experimental design, drafted the paper, and procured funding for all aspects of the work. Behavioral testing and data collection were performed by H. Dollish, S. Kaladchibachi, and D. Negelspach. All the authors contributed to discussing and interpreting the findings and approved the final version of the manuscript for submission.

---

**Table 1.** Seasonal PRCs to light: Summary of phase-shifting data in a high winter-like photoschedule. Response after a 15-min broadspectrum pulse.

| Zeitgeber Time | Photoperiod<br>(Composition of Light-Dark Cycle) | $\Delta$ Phase Shift, h<br>(*n*) |
| --- | --- | --- |
| Lights-on<br>0 | 8 hours of light • 16 hours of darkness | -0.04 ± 0.13 (32) |
| 1 | 8 hours of light • 16 hours of darkness | -0.06 ± 0.09 (32) |
| 2 | 8 hours of light • 16 hours of darkness | -0.11 ± 0.11 (32) |
| 3 | 8 hours of light • 16 hours of darkness | -0.13 ± 0.11 (31) |
| 4 | 8 hours of light • 16 hours of darkness | -0.04 ± 0.10 (32) |
| 5 | 8 hours of light • 16 hours of darkness | -0.00 ± 0.10 (32) |
| 6 | 8 hours of light • 16 hours of darkness | -0.05 ± 0.08 (32) |
| 7 | 8 hours of light • 16 hours of darkness | -0.40 ± 0.08 (32) |
| 8<br>Lights-off | 8 hours of light • 16 hours of darkness | -0.47 ± 0.07 (32) |
| 9 | 8 hours of light • 16 hours of darkness | -0.60 ± 0.09 (32) |
| 10 | 8 hours of light • 16 hours of darkness | -2.54 ± 0.12 (32) |
| 11 | 8 hours of light • 16 hours of darkness | -4.49 ± 0.17 (31) |
| 12 | 8 hours of light • 16 hours of darkness | -4.25 ± 0.14 (32) |
| 13 | 8 hours of light • 16 hours of darkness | -2.75 ± 0.16 (32) |
| 14 | 8 hours of light • 16 hours of darkness | 0.24 ± 0.18 (30) |
| 15 | 8 hours of light • 16 hours of darkness | 1.78 ± 0.16 (31) |
| 16 | 8 hours of light • 16 hours of darkness | 2.76 ± 0.16 (32) |
| 17 | 8 hours of light • 16 hours of darkness | 3.17 ± 0.15 (32) |
| 18 | 8 hours of light • 16 hours of darkness | 2.88 ± 0.15 (32) |
| 19 | 8 hours of light • 16 hours of darkness | 2.72 ± 0.12 (31) |
| 20 | 8 hours of light • 16 hours of darkness | 1.85 ± 0.12 (32) |
| 21 | 8 hours of light • 16 hours of darkness | 1.34 ± 0.14 (32) |
| 22 | 8 hours of light • 16 hours of darkness | 0.83 ± 0.12 (32) |
| 23 | 8 hours of light • 16 hours of darkness | 0.24 ± 0.15 (31) |

**Table 2.** Seasonal PRCs to light: Summary of phase-shifting data in a winter-like photoschedule. Response after a 15-min broadspectrum pulse.

| Zeitgeber Time | Photoperiod<br>(Composition of Light-Dark Cycle) | $\Delta$ Phase Shift, h<br>(n) |
| --- | --- | --- |
| Lights-on<br>0 | 10 hours of light • 14 hours of darkness | $1.17 \pm 0.12$ (32) |
| 1 | 10 hours of light • 14 hours of darkness | $0.78 \pm 0.11$ (32) |
| 2 | 10 hours of light • 14 hours of darkness | $0.67 \pm 0.13$ (26) |
| 3 | 10 hours of light • 14 hours of darkness | $0.26 \pm 0.12$ (32) |
| 4 | 10 hours of light • 14 hours of darkness | $0.15 \pm 0.11$ (32) |
| 5 | 10 hours of light • 14 hours of darkness | $0.26 \pm 0.09$ (32) |
| 6 | 10 hours of light • 14 hours of darkness | $-0.06 \pm 0.10$ (32) |
| 7 | 10 hours of light • 14 hours of darkness | $0.11 \pm 0.09$ (32) |
| 8 | 10 hours of light • 14 hours of darkness | $-0.04 \pm 0.10$ (32) |
| 9 | 10 hours of light • 14 hours of darkness | $-0.15 \pm 0.09$ (32) |
| 10<br>Lights-off | 10 hours of light • 14 hours of darkness | $-0.29 \pm 0.08$ (32) |
| 11 | 10 hours of light • 14 hours of darkness | $-1.25 \pm 0.12$ (32) |
| 12 | 10 hours of light • 14 hours of darkness | $-3.77 \pm 0.16$ (32) |
| 13 | 10 hours of light • 14 hours of darkness | $-4.01 \pm 0.15$ (32) |
| 14 | 10 hours of light • 14 hours of darkness | $-3.15 \pm 0.14$ (32) |
| 15 | 10 hours of light • 14 hours of darkness | $-1.11 \pm 0.29$ (32) |
| 16 | 10 hours of light • 14 hours of darkness | $0.97 \pm 0.22$ (31) |
| 17 | 10 hours of light • 14 hours of darkness | $2.14 \pm 0.14$ (31) |
| 18 | 10 hours of light • 14 hours of darkness | $2.82 \pm 0.11$ (31) |
| 19 | 10 hours of light • 14 hours of darkness | $2.99 \pm 0.13$ (31) |
| 20 | 10 hours of light • 14 hours of darkness | $2.67 \pm 0.13$ (30) |
| 21 | 10 hours of light • 14 hours of darkness | $2.64 \pm 0.13$ (29) |
| 22 | 10 hours of light • 14 hours of darkness | $1.93 \pm 0.13$ (32) |
| 23 | 10 hours of light • 14 hours of darkness | $1.64 \pm 0.09$ (32) |

**Table 3.**
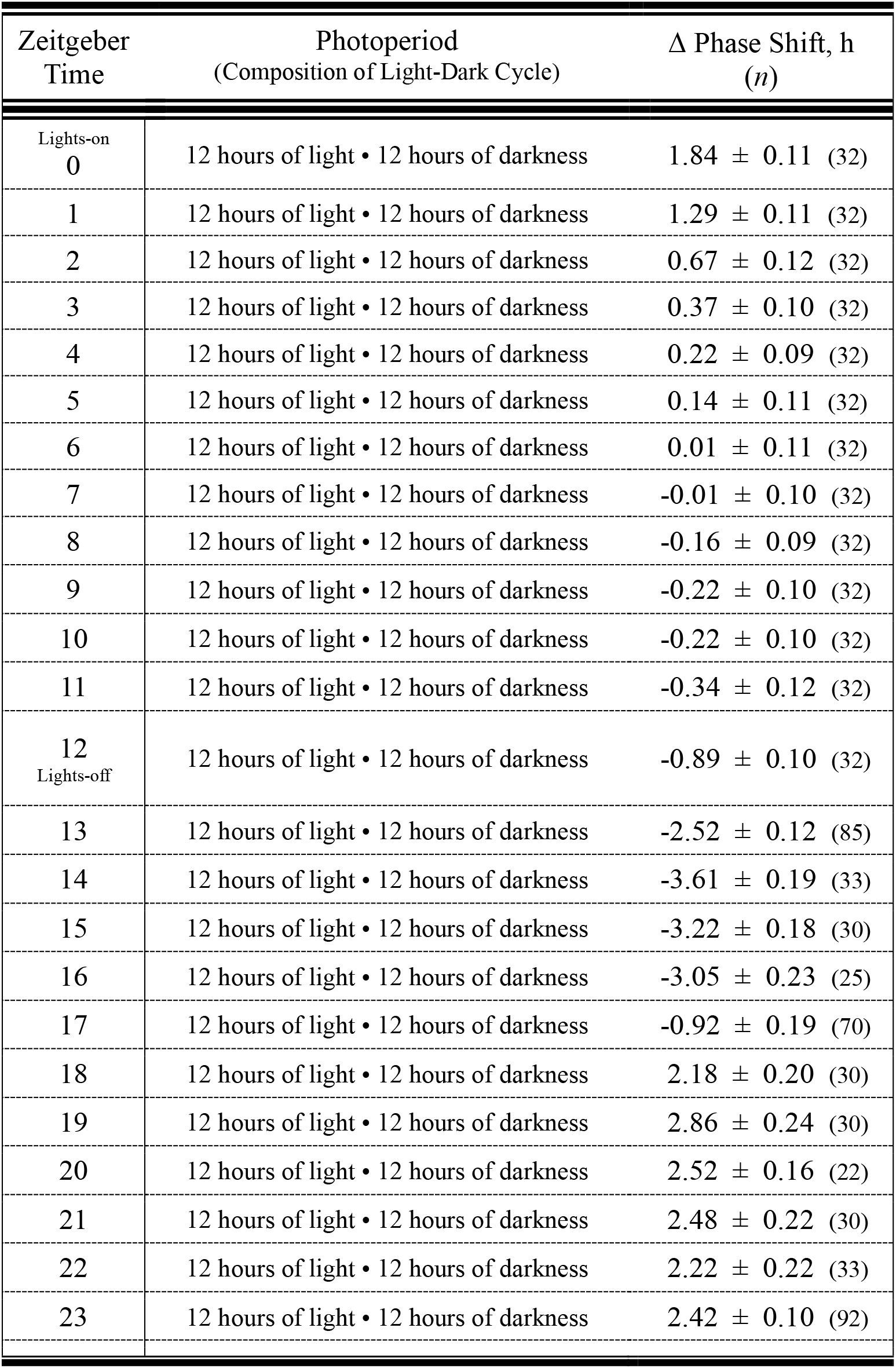
Seasonal PRCs to light: Summary of phase-shifting data within an equinoctial photoschedule. Response after a 15-min broadspectrum pulse.

| Zeitgeber Time | Photoperiod<br>(Composition of Light-Dark Cycle) | $\Delta$ Phase Shift, h<br>(*n*) |
| --- | --- | --- |
| Lights-on<br>0 | 12 hours of light • 12 hours of darkness | $1.84 \pm 0.11$ (32) |
| 1 | 12 hours of light • 12 hours of darkness | $1.29 \pm 0.11$ (32) |
| 2 | 12 hours of light • 12 hours of darkness | $0.67 \pm 0.12$ (32) |
| 3 | 12 hours of light • 12 hours of darkness | $0.37 \pm 0.10$ (32) |
| 4 | 12 hours of light • 12 hours of darkness | $0.22 \pm 0.09$ (32) |
| 5 | 12 hours of light • 12 hours of darkness | $0.14 \pm 0.11$ (32) |
| 6 | 12 hours of light • 12 hours of darkness | $0.01 \pm 0.11$ (32) |
| 7 | 12 hours of light • 12 hours of darkness | $-0.01 \pm 0.10$ (32) |
| 8 | 12 hours of light • 12 hours of darkness | $-0.16 \pm 0.09$ (32) |
| 9 | 12 hours of light • 12 hours of darkness | $-0.22 \pm 0.10$ (32) |
| 10 | 12 hours of light • 12 hours of darkness | $-0.22 \pm 0.10$ (32) |
| 11 | 12 hours of light • 12 hours of darkness | $-0.34 \pm 0.12$ (32) |
| 12<br>Lights-off | 12 hours of light • 12 hours of darkness | $-0.89 \pm 0.10$ (32) |
| 13 | 12 hours of light • 12 hours of darkness | $-2.52 \pm 0.12$ (85) |
| 14 | 12 hours of light • 12 hours of darkness | $-3.61 \pm 0.19$ (33) |
| 15 | 12 hours of light • 12 hours of darkness | $-3.22 \pm 0.18$ (30) |
| 16 | 12 hours of light • 12 hours of darkness | $-3.05 \pm 0.23$ (25) |
| 17 | 12 hours of light • 12 hours of darkness | $-0.92 \pm 0.19$ (70) |
| 18 | 12 hours of light • 12 hours of darkness | $2.18 \pm 0.20$ (30) |
| 19 | 12 hours of light • 12 hours of darkness | $2.86 \pm 0.24$ (30) |
| 20 | 12 hours of light • 12 hours of darkness | $2.52 \pm 0.16$ (22) |
| 21 | 12 hours of light • 12 hours of darkness | $2.48 \pm 0.22$ (30) |
| 22 | 12 hours of light • 12 hours of darkness | $2.22 \pm 0.22$ (33) |
| 23 | 12 hours of light • 12 hours of darkness | $2.42 \pm 0.10$ (92) |

**Table 4.** Seasonal PRCs to light: Summary of phase-shifting data in a summer-like photoschedule. Response after a 15-min broadspectrum pulse.

| Zeitgeber Time | Photoperiod<br>(Composition of Light-Dark Cycle) | $\Delta$ Phase Shift, h<br>(*n*) |
| --- | --- | --- |
| Lights-on<br>0 | 14 hours of light • 10 hours of darkness | $2.88 \pm 0.10$ (32) |
| 1 | 14 hours of light • 10 hours of darkness | $2.36 \pm 0.11$ (32) |
| 2 | 14 hours of light • 10 hours of darkness | $1.93 \pm 0.09$ (32) |
| 3 | 14 hours of light • 10 hours of darkness | $1.43 \pm 0.14$ (32) |
| 4 | 14 hours of light • 10 hours of darkness | $1.06 \pm 0.09$ (32) |
| 5 | 14 hours of light • 10 hours of darkness | $0.75 \pm 0.10$ (32) |
| 6 | 14 hours of light • 10 hours of darkness | $0.34 \pm 0.11$ (32) |
| 7 | 14 hours of light • 10 hours of darkness | $0.14 \pm 0.08$ (32) |
| 8 | 14 hours of light • 10 hours of darkness | $0.06 \pm 0.09$ (31) |
| 9 | 14 hours of light • 10 hours of darkness | $0.10 \pm 0.09$ (32) |
| 10 | 14 hours of light • 10 hours of darkness | $-0.02 \pm 0.10$ (32) |
| 11 | 14 hours of light • 10 hours of darkness | $-0.17 \pm 0.12$ (32) |
| 12 | 14 hours of light • 10 hours of darkness | $-0.20 \pm 0.08$ (32) |
| 13 | 14 hours of light • 10 hours of darkness | $-0.56 \pm 0.11$ (32) |
| 14<br>Lights-off | 14 hours of light • 10 hours of darkness | $-0.78 \pm 0.14$ (31) |
| 15 | 14 hours of light • 10 hours of darkness | $-1.72 \pm 0.17$ (32) |
| 16 | 14 hours of light • 10 hours of darkness | $-2.73 \pm 0.14$ (32) |
| 17 | 14 hours of light • 10 hours of darkness | $-4.32 \pm 0.14$ (31) |
| 18 | 14 hours of light • 10 hours of darkness | $-4.57 \pm 0.16$ (31) |
| 19 | 14 hours of light • 10 hours of darkness | $-2.78 \pm 0.22$ (32) |
| 20 | 14 hours of light • 10 hours of darkness | $1.51 \pm 0.17$ (32) |
| 21 | 14 hours of light • 10 hours of darkness | $2.84 \pm 0.11$ (32) |
| 22 | 14 hours of light • 10 hours of darkness | $3.40 \pm 0.14$ (31) |
| 23 | 14 hours of light • 10 hours of darkness | $3.01 \pm 0.13$ (32) |

**Table 5.**
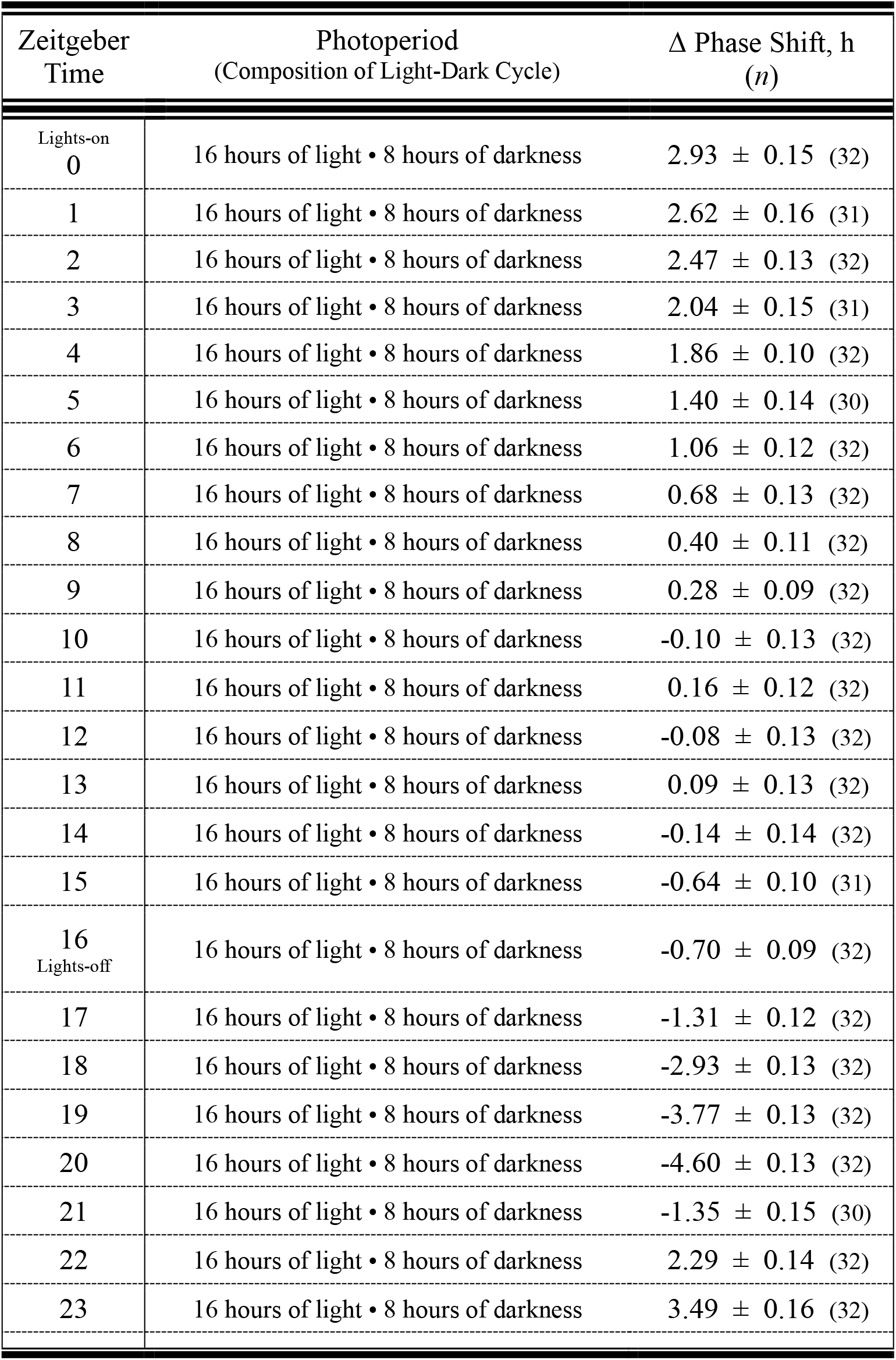
Seasonal PRCs to light: Summary of phase-shifting data in a high summer-like photoschedule. Response after a 15-min broadspectrum pulse.

